# Similar deamination activity but different phenotypic outcomes induced by APOBEC3 enzymes in breast epithelial cells

**DOI:** 10.1101/2023.03.29.534844

**Authors:** Milaid Granadillo Rodríguez, Lai Wong, Linda Chelico

**Author notes:** Correspondence: Linda Chelico.

## Abstract

APOBEC3 (A3) enzymes deaminate cytosine to uracil in viral single-stranded DNA as a mutagenic barrier for some viruses. A3-induced deaminations can also occur in human genomes resulting in an endogenous source of somatic mutations in multiple cancers. However, the roles of each A3 are unclear since few studies have assessed these enzymes in parallel. Thus, we developed stable cell lines expressing A3A, A3B, or A3H Hap I using non-tumorigenic MCF10A and tumorigenic MCF7 breast epithelial cells, to assess their mutagenic potential and cancer phenotypes in breast cells. The activity of these enzymes was characterized by γH2AX foci formation and *in vitro* deamination. Cell migration, and soft agar colony formation assays assessed cellular transformation potential. We found that all three A3 enzymes had similar γH2AX foci formation, despite different deamination activity *in vitro*. Notably, in nuclear lysates the *in vitro* deaminase activity of A3A, A3B, and A3H did not require digestion of cellular RNA, in contrast to A3B and A3H in whole cell lysates. Their similar activities in cells nonetheless resulted in distinct phenotypes where A3A decreased colony formation in soft agar, A3B decreased colony formation in soft agar after hydroxyurea treatment, and A3H Hap I promoted cell migration. Overall, we show that *in vitro* deamination data does not always reflect in cell deamination, all three A3s induce somatic mutagenesis, and the impact of each is different.

## 1 Introduction

The human APOBEC3 (A3) enzymes belong to the AID/APOBEC family and consist of seven members (A3A, A3B, A3C, A3D, A3F, A3G, and A3H) that deaminate cytosine to form uracil on single stranded (ss)DNA (Refsland and Harris, 2013). Since uracil templates for the addition of adenosine or initiates base excision repair (BER) to remove the uracil, these enzymes are promutagenic. Normally this is used as an intrinsic barrier to retroviruses, endogenous retroelements and some DNA viruses where A3-induced uracils in these genomes causes their inactivation or degradation (Arias et al., 2012;Feng et al., 2014;Cheng et al., 2021). Accordingly, A3 enzymes are normally only highly expressed in germ cells, lymphoid cells, and epithelial cells in response to viral infection (Jarmuz et al., 2002;Koning et al., 2009;Okeoma et al., 2010;Refsland et al., 2010;Milewska et al., 2018;Wei et al., 2020). However, some AID/APOBEC enzymes have also been harnessed as tools in genome editing (Evanoff and Komor, 2019). Acting as cytosine base editors (CBEs) by linking an AID/APOBEC enzyme to catalytically inactive CRISPR-Cas9 the formation of uracil by AID/APOBEC catalysis can be used to induce C→T mutations in somatic cells (Komor et al., 2016;St Martin et al., 2018;McGrath et al., 2019;Liu et al., 2020;Zhao et al., 2021). However, their off-target effects need to be controlled. This is especially important since A3 enzymes are a source of somatic mutations in at least 70% of cancer types with 25-50% of sequenced tumor genomes containing A3-induced mutations (Petljak and Alexandrov, 2016;Alexandrov et al., 2020;Bergstrom et al., 2022). Understanding the natural mutagenic potential of A3 enzymes in the genome, can inform research on their roles as CBEs and our understanding of the origins of somatic mutations in tumor cells.

To induce mutations in genomic DNA, it is of course important that the enzymes localize to the nucleus, but the enzymes must also be able to compete for ssDNA with Replication Protein A (RPA) during dynamic and transient processes such as replication, transcription, and DNA repair (Swanton et al., 2015;Adolph et al., 2017b;Mertz et al., 2017;Adolph et al., 2018;Green and Weitzman, 2019;Wong et al., 2021). Only A3A, A3B, A3C, and A3H can enter the nucleus (Vieira and Soares, 2013). A3G can enter the nucleus in certain cancer cells and after ionizing radiation treatment (Talluri et al., 2021;Britan-Rosich et al., 2022;Liu et al., 2023). A3H has seven major haplotypes and although all can enter the nucleus to varying degrees, A3H Haplotype I (Hap I) is the only one that is predominantly nuclear (Li and Emerman, 2011;Wang et al., 2011). A3B has a nuclear localization signal (NLS) and A3A and A3C enter the nucleus by diffusion (Bogerd et al., 2006;Chen et al., 2006;Lackey et al., 2012).

To date, only A3A, A3B, and A3H Hap I have been implicated in processes related to somatic mutagenesis and cancer using retrospective data from The Cancer Genome Atlas (TCGA) or other methods where whole genome sequencing and mRNA expression can be analyzed (Burns et al., 2013a;Middlebrooks et al., 2016;Starrett et al., 2016;Yan et al., 2016;Glaser et al., 2018;Chen et al., 2019;Roper et al., 2019;Hix et al., 2020;Law et al., 2020;Chervova et al., 2021;Petljak et al., 2022). Suppression of A3 mRNA transcription provides the main protective measure against somatic mutagenesis, although A3B and A3A transcription is upregulated in response to DNA replication stress, an early sign of cellular transformation and viral infection (Kanu et al., 2016;Oh et al., 2021). A3A activity also appears to be regulated by binding partners that suppress catalytic activity (Aynaud et al., 2012;Green et al., 2021). A3H Hap I is ubiquitinated in cells and the resulting short half-life decreases its catalytic activity (Chesarino and Emerman, 2020). A3B and A3H Hap I cytidine deaminase activity can also be suppressed by binding cellular RNA, but the implications of this for their activity in the nucleus is not known (Cortez et al., 2019).

The formation of promutagenic uracils through the intrinsic deaminase activity of these enzymes occurs in a sequence-specific manner and those involved in cancer preferentially deaminate the central cytosine in 5’TCW trinucleotide motifs (where W = A or T on ssDNA) to form uracil (Refsland and Harris, 2013). Uracil-DNA glycosylase (UDG) is a highly conserved repair enzyme that initiates BER and catalyzes the excision of uracil from ssDNA and double-stranded (ds)DNA and prevents mutagenesis from A3 enzymes (Friedberg et al., 2005). In this context, the uracil DNA glycosylase inhibitor (UGI) protein from bacteriophage PBS2 has been a useful tool to determine if A3-catalyzed uracils induce DNA damage and mutagenesis (Richardson et al., 2014). Some uracils escape BER and result in C→T mutations when uracil is used as a template during replication or an error prone process mediated by Rev1 can result in C→G mutations due to insertion of C opposite the abasic site created during BER (Nik-Zainal et al., 2012;Roberts et al., 2012;Chan et al., 2013;Henderson and Fenton, 2015). These two types of mutations form the basis of APOBEC-induced single base substitution (SBS) signatures SBS2 (C→T) and SBS13 (C→G) in cancer genomes (Alexandrov et al., 2013).

A3B was first identified as a significant mutational source in breast cancer (BC), but was later identified in other cancer types including bladder, cervical, lung, and head-and-neck cancers where it was evident that there was a positive correlation between A3B expression and the total APOBEC-mutation load (Burns et al., 2013a;Burns et al., 2013b). A3B is overexpressed in almost 50% of BC, the majority of BC cell lines, and was linked to DNA deaminase activity in BC cell extracts (Burns et al., 2013a;Petljak et al., 2022). High A3B expression levels have been associated with poor clinical outcomes for estrogen receptor-positive BC and can promote tamoxifen resistance (Sieuwerts et al., 2014;Law et al., 2016). However, the frequency of A3B mutations is not always correlated with A3B expression levels (Nik-Zainal et al., 2014;Roberts and Gordenin, 2014;Cescon et al., 2015;Petljak et al., 2022). In addition, 22% of humans have a deletion polymorphism resulting in loss of the A3B gene and sequencing of BC tumors from individuals carrying this deletion indicated that these tumors contained APOBEC-induced mutations at higher abundance than the non-carriers, consistent with recent data in cell lines with artificial A3A or A3B deletions introduced (Kidd et al., 2007;Nik-Zainal et al., 2014;Petljak et al., 2022). Although A3A mRNA levels in BC are much lower than A3B, they are more significantly correlated with BC and A3A appears to be more active than A3B *in vitro*, suggesting that less steady state A3A is needed in cells to induce somatic mutagenesis (Cortez et al., 2019;Petljak et al., 2022). Another study found the A3A mutational footprints in tumors, but no corresponding A3A expression and suggested that A3A is upregulated early, but later inactivated, perhaps due to being the most active deaminase that could cause cell death through its activity over time, resulting in episodic expression (Landry et al., 2011;Mussil et al., 2013;Chan et al., 2015). In addition, one study of APOBEC-induced mutations from A3B deleted BC tumors revealed that the only tumors displaying the APOBEC mutation signature also contained at least one allele of A3H Hap I, providing correlative evidence that this enzyme may be the additional source of mutagenesis (Starrett et al., 2016).

The previous studies are not conclusive about the relative contribution of A3A, A3B, and A3H Hap I to BC mutagenesis. A major issue has been the use of retrospective patient samples (Petljak et al., 2019), reliance on mRNA as an indicator of enzyme activity (Jalili et al., 2020), or a lack of studies where A3 enzymes were tested in parallel. This has led to questions as to whether APOBEC-induced mutations are drivers or passengers in the overall tumorigenesis process. Most of the previous A3A research was done in yeast, cell lines and *in silico*, but recently there has been direct evidence of A3A tumorigenic potential *in vivo* (Chan et al., 2015;Glaser et al., 2018;Cortez et al., 2019;Jalili et al., 2020;Law et al., 2020). The mutation signature of A3A is also higher than A3B in MDA-MB-453 and BT474 breast carcinoma cell lines (Petljak et al., 2022). Direct comparison of deamination activity of A3A and A3B in whole cell (WC) lysates of the breast carcinoma cell lines BT474, CAMA-1, and MDA-MB-453 showed that A3A was more active than A3B, but part of the reason for higher activity of A3A is that it was not inhibited by RNA, unlike A3B and A3H Hap I (Cortez et al., 2019).

Although activities of the A3 enzymes have been characterized in various ways, there is still not a comparison of deamination activity in nuclear lysates, where there may be less inhibitory RNA than WC lysates. In addition, there have been no studies determining if A3- induced mutations cause any specific cancer cell phenotype. Here, we aimed to compare in parallel A3A, A3B, and A3H Hap I. We chose BC as a model because all three of the A3s have been implicated (Burns et al., 2013a;Burns et al., 2013b;Chan et al., 2015;Starrett et al., 2016;Cortez et al., 2019). Using two human breast epithelial cell lines, MCF10A (non-tumorigenic, immortalized mammary epithelial cell line) and MCF7 (tumorigenic) that have both borderline undetectable endogenous A3 expression (Burns et al., 2013a), we developed stably transduced cell lines expressing either Flag-tagged A3A, A3B or A3H Hap I and tested the induced DNA damage, deamination activity, and ability to induce cancer cell phenotypes.

## 2 Materials and methods

### 2.1 Cell culture and Generation of Stable Cell Lines Expressing A3s

The 293T, HCC1428, MCF10A, and MCF7 cell lines were obtained from ATCC and cultured as recommended by ATCC, excluding antibiotics. Lentiviruses were generated by transfection of a pLVX-3x Flag vector with psPAX2 and pMD2.G into 293T cells. The transduction of MCF10A and MCF7 with the resulting lentiviruses expressing Flag tagged - A3A, -A3B, and -A3H Hap I was performed for 16 h in medium containing 8 µg/mL polybrene. Transduced MCF10A and MCF7 cells were selected with 1 µg/mL and 2 µg/mL puromycin, respectively, for 7 d and maintained as polyclonal cell lines with 0.25 µg/mL puromycin.

### 2.2 Immunoblotting

The MCF10A- and MCF7- derived stable cell lines were incubated for 72 h with doxycycline (dox) concentrations ranging from 0.05 to 2 µg/mL to induce the expression of A3-Flag enzymes. Cells were lysed with 2X Laemmli buffer and WC, cytoplasmic (Cyt), and nuclear (Nuc) lysates obtained after fractionation were resolved by SDS-PAGE. Antibodies used are labeled on each immunoblot. Secondary detection was performed using Licor IRDye antibodies produced in donkey (IRDye 680-labeled anti-rabbit and IRDye 800-labeled anti-mouse). Quantification of band intensities was performed using Odyssey Software with normalization of each experimental lane to its respective α-tubulin, which was detected in parallel on the same blot. The antibodies purchased from Sigma-Aldrich were: Anti-α-Tubulin antibody, Mouse monoclonal (T8203/clone AA13, purified from hybridoma cell culture, Public ID AB_1841230, used at 1:1000); Anti-alpha-Tubulin antibody, Rabbit monoclonal (SAB5600206/clone RM113, used at 1:5000); ANTI-FLAG® antibody produced in rabbit (F7425, Public ID AB_439687, used at 1:1000); Monoclonal ANTI-FLAG® M2 antibody produced in mouse (F1804/Clone M2, Public ID AB_262044, used at 1:1000); and Anti-HA antibody produced in rabbit (H6908, Public ID AB_260070, used at 1:1000). The antibody purchased from Thermo Fisher Scientific was Histone H2B monoclonal antibody (MA5- 31566/Clone GT387, Public ID AB_2787193, used at 1:500). The antibodies purchased from LI-COR Biosciences were IRDye 680RD Donkey anti-Rabbit IgG, Secondary antibody (926- 68073, Public ID AB_2895657, used at 1:10,000) and IRDye® 800CW Donkey anti-Mouse IgG Secondary Antibody (926-32212, Public ID AB_621847, used at 1:10,000).

### 2.3 qRT-PCR

The RNA preparation, cDNA synthesis and qPCR were carried out according to Refsland et al. (Refsland et al., 2010). MCF10A and MCF7 parental and derived cell lines were either untreated or treated with dox for 72 h to induce the expression of A3-Flag enzymes or as a mock control. The primers used for A3B, A3C, A3G, and TBP were as reported in Refsland et al. (Refsland et al., 2010). The primers for A3A and A3H are listed in Table S1.

### 2.4 Immunofluorescence Microscopy

MCF10A- and MCF7- derived stable and transduced cell lines were grown on glass slides and treated for 24 and 72 h with dox for the expression of the A3-Flag enzymes and fixed with 100% cold methanol for 20 min. Where indicated, the cells were transfected with an expression vector encoding the bacteriophage PBS2 UGI protein 24 h before the dox induction (Feng et al., 2017). Also where indicated, the cells were transduced with catalytic mutants of each A3 enzyme, A3A E72Q, A3B E255Q, or A3H Hap I E56Q. Catalytic mutants were generated by site directed mutagenesis of the pLVX-3x Flag A3 vectors (Wang and Malcolm, 1999). Primers are listed in Table S2. After dox treatment, the cells were permeabilized and stained as previously described (Hix et al., 2020). Quantification of cells with γH2AX foci is based on counting of at least 100 cells. The antibodies purchased from Sigma-Aldrich were: Anti-phospho-H2AFX (pSer139) antibody produced in rabbit (SAB4300213, Public ID 10620164, used at 1:200) and Monoclonal ANTI-FLAG® M2 antibody produced in mouse (F1804/Clone M2, Public ID AB_262044, used at 1:200).

### 2.5 Preparation of Whole Cell Lysates and Cell Fractionation

The 293T cells were transfected using GeneJuice (EMD Millipore) with 1 µg of pcDNA, pcDNA-A3A-3xHA, and pVIVO-A3H Hap I-HA for 40 h or 2 µg of pcDNA-A3B- 3xHA for 48 h to obtain similar protein levels. Additionally, 293T cells that were not transfected were processed to obtain Nuc lysates. MCF10A- and MCF7- derived stable cell lines were treated with dox for 72 h. Cells were harvested and either lysed using sonication in HED buffer (25 mM HEPES, 10% glycerol, 1 mM DTT, and EDTA-free protease inhibitor (Roche, Basel, Switzerland)) containing 0.1, 0.5 or 5 mM EDTA or 2X Laemmli buffer to get the WC lysates for deamination or immunoblot, respectively, or were fractionated. For fractionation, cells were washed with 1X PBS and pelleted by centrifugation at 500 x *g* for 10 min. The cell pellet was resuspended in Cyt extraction buffer (10 mM HEPES pH 7.4, 10 mM KCl, 1.5 mM MgCl2, 340 mM sucrose, 0.5 mM EDTA, 1 mM DTT, and EDTA-free protease inhibitor (Roche)). After 10 min of incubation on ice, 12.5 µL of 10% NP-40 per 1 x 10^6^ cells was added and followed by vortex mixing at high speed for 15 s. The samples were centrifuged at 3 300 x *g* for 10 min and the supernatant was saved as Cyt lysate. The remaining pellet was washed once with Cyt extraction buffer and lysed with Nuc extraction buffer (50 mM HEPES pH 7.4, 500 mM NaCl, 1.5 mM MgCl2, 0.1% NP 40, 0.5 mM EDTA, 1 mM DTT, and EDTA-free protease inhibitor (Roche)) for 30 min at 4°C. After centrifugation at 13 000 x *g*, the resulting supernatant was saved as Nuc lysate.

### 2.6 Protein purification

The A3A, A3B, and A3H Hap VII were purified from *Sf*9 cells infected with a recombinant baculovirus expressing a GST-tagged A3. Purification and GST-tag cleavage has been previously described (Love et al., 2012;Feng et al., 2015;Adolph et al., 2017b).

### 2.7 Deamination assays

#### 2.7.1 Deamination activity of cell expressed enzymes

The 293T cells were either transfected or not with pcDNA or pVIVO A3 expression plasmids, as described for preparation of WC lysates, for 40-48 h and MCF10A-derived stable cell lines were treated with dox for 72 h. Deamination assays were performed using a protocol adapted from Serebrenik et al. (Serebrenik et al., 2019). For either non-transfected or transfected 293T cells, the RNase A (6 µg/mL, Roche) was added or not directly to the reaction mixture. Prior to performing the deaminase assay for the MCF10A stable cell lines, WC lysates were either pre-treated or not with RNase A (100 μg/mL) for 15 min at 37°C. ). Deamination reactions were processed, resolved and analyzed as previously described (Adolph et al., 2017a;Serebrenik et al., 2019).

#### 2.7.2 Deamination activity of purified enzymes in Nuc lysates

For the Nuc lysates from the non-transfected 293T cells, the purified enzymes A3A, A3B, or A3H Hap VII were added to the reactions together with 40 U of murine RNase inhibitor (NEB, Ipswich, Massachusetts). Activity of cell lysates was evaluated in reactions containing 43 nt or 85 nt linear or 21 nt hairpin substrates (Table S3). Deamination reactions were processed, resolved and analyzed as previously described (Adolph et al., 2017a;Serebrenik et al., 2019).

### 2.8 Soft agar assay

MCF10A and MCF7 parental cell lines and cell lines transduced to express Flag-tagged A3s were either untreated or treated for 24 h with 0.05 to 2 μg/mL dox. To examine the effect of hydroxyurea (HU) on anchorage independent growth, MCF10A and MCF7 (parental and A3-expressing cells) were either untreated or treated for 6 h with 2 mM or 12 mM HU, respectively, to stall replication (Supplementary Figures S1 and S2). A cell suspension consisting of 30 000 cells in 0.36% top agar (2x media (DMEM or EMEM), FBS, 4x MEM vitamin solution) was overlaid in duplicate in 6-well plates that contained 0.61% bottom agar (2x media, FBS, 4x MEM vitamin solution). Plates were maintained for 3 weeks and topped with media containing dox every 2 d to maintain A3 expression and prevent desiccation. After weeks, colonies were stained with crystal violet, visualized under a microscope, and quantified manually. The experiment was repeated three times and for each biological replicate, 10 fields were blindly counted.

### 2.9 Cell migration assay

The effect of A3 expression on the cell migration, in the presence or absence of HU, was evaluated using the QCM™ 24-well colorimetric cell migration assay (Millipore, Burlington, Massachusetts). MCF10A- and MCF7- derived stable cell lines were either untreated, treated with dox, or treated with dox+HU. The cells were treated with HU twice, at days 3 and 6 and at HU concentrations of 2 mM and 12 mM for MCF10A and MCF7 stable cell lines, respectively. Cells were starved for 16 h prior to the assay in media containing 0.5% serum and then seeded (3 × 10^5^ cells) onto the upper chamber in media containing 0.5% serum. Media containing either 0.5% or 10% serum was loaded into the lower chambers. Migration during 48 h toward the lower chamber was determined by colorimetric measurement at 560 nm.

### 2.10 Statistical analysis

The results are presented as mean ± standard deviation (SD). Two-tailed t test was applied for comparisons of two groups, while one-way analysis of variance (ANOVA) followed by Holm-Sidak post hoc test was used for multiple groups. The results were considered statistically significant at a p value of <0.05. Statistical analyses were performed using SigmaPlot 11.0 software.

## 3 Results

### 3.1 A3A, A3B, and A3H Hap I protein expression levels in the stable cell lines

In order to compare the cellular activities of A3A, A3B, and A3H Hap I, we first generated polyclonal MCF10A and MCF7 inducible cell lines to stably express the Flag-tagged A3s (Figure 1A). The cells were then treated with different dox concentrations ranging from 0.05 to 2 µg/mL to determine the concentration that would induce similar protein expression levels (Figure 1B-C). We were able to obtain similar protein expression for A3A and A3B in MCF10A and MCF7 cells, whereas A3H Hap I expression was slightly higher, even with using lower dox induction levels. This was determined by calculating the relative protein levels after normalizing each Flag band to the α-tubulin in the same lane. The MCF10A- and MCF7- relative protein levels are comparable within each set but not to each other since they were detected on different immunoblots (Supplementary Figure S3). Using these conditions that had equal A3-Flag expression levels, the mRNA expression levels were assessed by qRT-PCR to determine if there was any expression of endogenous A3 enzymes that can access the nucleus (Figure 1D-E). For the parental, non-transduced MCF10A and MCF7 cell lines, endogenous *A3A* mRNA was not detected, but there was basal endogenous mRNA expression detected for *A3B* and *A3H Hap I* after treatment with dox (Figure 1D-E, MCF10A or MCF7 on x-axis). After dox induction in transduced cells (labeled A3A, A3B, or A3H Hap I on x-axis), the level of *A3A* mRNA for both MCF10A and MCF7 cells was low, showing only 0.75- and 1.42- fold increases, respectively, although the steady state protein expression was similar to A3B (Figure 1B-E). In contrast, *A3B* mRNA in both transduced cell lines was induced over 60-fold (Figure 1D-E). *A3H Hap I* mRNA induction in transduced cell lines was over 100-fold in MCF10A cells and 50-fold in MCF7 cells (Figure 1D-E). We also determined if A3C or A3G were expressed in MCF10A or MCF7 non-transduced cells since they are the only other A3 enzymes with access to the nucleus (Lackey et al., 2013;Liu et al., 2023). However, we did not detect any *A3C* or *A3G* mRNA (data not shown).

**Figure 1.**
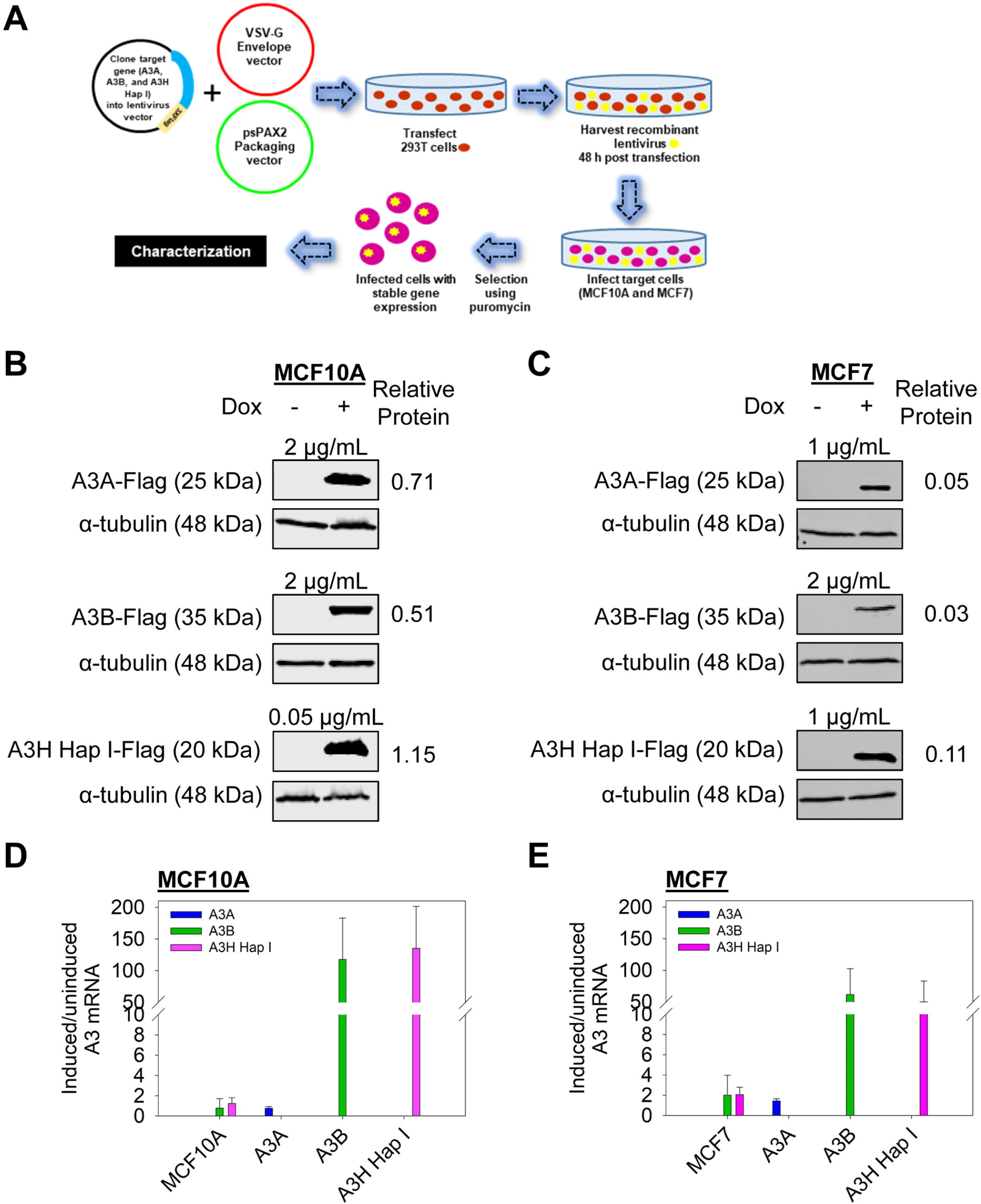
A3 protein and mRNA transcript levels in the stable cell lines. A. Schematic of the generation of the stable cell lines derived from MCF10A and MCF7 cells. **B-C.** Expression of A3A, A3B, and A3H Hap I in MCF10A and MCF7 cells. The cells were either uninduced or dox induced for 72 h and the protein expression was assessed by immunoblot. To obtain similar protein expression levels the following dox concentrations were used for each cell line: MCF10A A3A (2 µg/mL dox); MCF10A A3B (2 µg/mL dox); MCF10A A3H Hap I (0.05 µg/mL dox); MCF7 A3A (1 µg/mL dox); MCF7 A3B (2 µg/mL dox); MCF7 A3H Hap I (1 µg/mL dox). Three biologically independent experiments were conducted. Approximate molecular weight (kDa) of each protein is shown based on the closest molecular weight marker band (Supplementary Figure S3). Bands were quantified from one representative experiment and the Flag intensity was divided by the α-tubulin intensity to account for potential loading differences. Representative results are shown as relative protein levels. Blots were cropped to improve conciseness of presentation and uncropped blots are shown in Supplementary Figure S3. **D-E.** The mRNA expression measured by qRT-PCR is shown as the fold change of each *A3* for induced compared to uninduced conditions after normalization of each to *TBP* expression. Error bars indicate the SD from the mean for three biologically independent replicates.

### 3.2 A3A, A3B, and A3H Hap I localize to the nucleus and induce γH2AX foci formation

We assessed the A3 localization and ability to induce DNA damage by immunofluorescence microscopy (Figure 2A). We included the non-transduced parental cells exposed to dox as the mock and also dox uninduced and induced conditions for the stable cell lines. For MCF10A- and MCF7- derived stable cell lines, immunofluorescence microscopy showed that A3A and A3H Hap I localized to the nucleus and cytoplasm, but A3B localized only to the nucleus (Figure 2B-G). To determine the γH2AX foci formation, which is a marker of stalled replication forks and dsDNA breaks, we analyzed cells by immunofluorescence microscopy 72 h post dox induction (Figure 2). All cell lines were also either transfected or not with a plasmid expressing UGI to demonstrate that the γH2AX foci were caused by removal of uracil by UDG. As a further control that the γH2AX foci were caused by A3-catalyzed deaminations, we also transduced cells with a catalytic mutant of A3A (E72Q), A3B (E255Q), or A3H Hap I (E56Q) (Figure 2B-G). The expression of A3A, A3B, and A3H Hap I in either MCF10A or MCF7 cells induced similar pan nuclear γH2AX staining that was blocked for A3- expressing cells that were also expressing UGI demonstrating that γH2AX foci formation was due to formation of A3-catalyzed uracils (Figure 2B-G). There was no γH2AX foci for any of the cells expressing the catalytic mutants of the A3 enzymes (Figure 2B-G).

**Figure 2.**
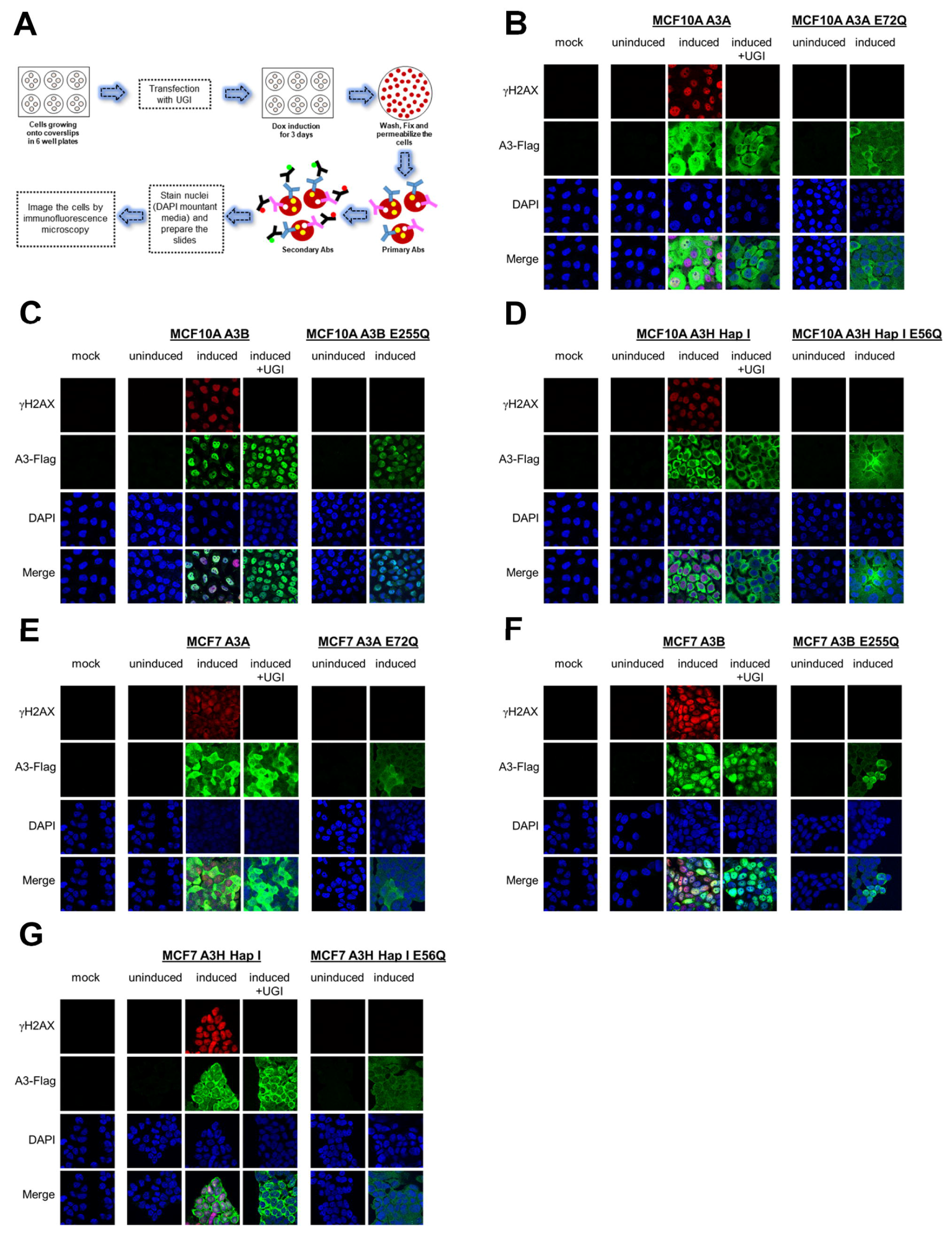
γH2AX foci formation by A3A, A3B, and A3H Hap I. A. Schematic representation of the protocol followed to assess the protein localization and γH2AX foci formation by immunofluorescence microscopy using antibodies (Abs). **B-D.** The MCF10A cell line was not transduced and treated with dox for 72 h (mock), transduced to express A3A, A3B, or A3H Hap I and not treated with dox for 72 h (uninduced), transduced to express A3A, A3B, or A3H Hap I and treated with dox for 72 h (induced) without or with transfection of a UGI expression plasmid (induced+UGI), transiently transduced with catalytic mutants of A3A (E72Q), A3B (E255Q), or A3H Hap I (E56Q) and not treated with dox for 72 h (uninduced), transiently transduced with catalytic mutants of A3A (E72Q), A3B (E255Q), or A3H Hap I (E56Q) and treated with dox for 72 h (induced). Green color identifies Flag-tagged A3 enzymes, red color indicates γH2AX foci formation, and blue color indicates nuclei staining. **E-G.** MCF7 parental cell line and MCF7-derived stable cell lines were subjected to same treatments as for MCF10A cell lines.

Previous studies have found different activities between A3A, A3B, and A3H Hap I, but these three enzymes have never been tested in parallel (Burns et al., 2013a;Burns et al., 2013b;Chan et al., 2015;Starrett et al., 2016;Cortez et al., 2019). Thus, to determine if there is any differences in the A3 activity in cells, we detected γH2AX foci after 24 h since the effect of the A3s appeared saturated at 72 h (Figure 2). After 24 h, we detected γH2AX foci formation in 47% of A3A-, 62% of A3B-, and 64% of A3H Hap I- expressing MCF10A cells (Figure 3A and Supplementary Figure S4). These data demonstrate that the formation of γH2AX foci from cytosine deamination is similar for all three A3 enzymes. For MCF7 cells, we detected γH2AX foci formation in 35% of A3A-, 51% of A3B-, 17% of A3H Hap I- expressing cells (Figure 3B and Supplementary Figure S4). For A3H Hap I in MCF7 cells the formation of γH2AX foci formation were ∼2-fold slower and detection at 72 h showed similar γH2AX foci to A3A and A3B after 24 h (56%, Figure 3B and Supplementary Figure S5). Altogether the results indicated that A3A, A3B, and A3H Hap I were all able to induce DNA damage in cells at equal amounts in MCF10A cells by 24 h and in MCF7 cells by 72 h.

**Figure 3.**
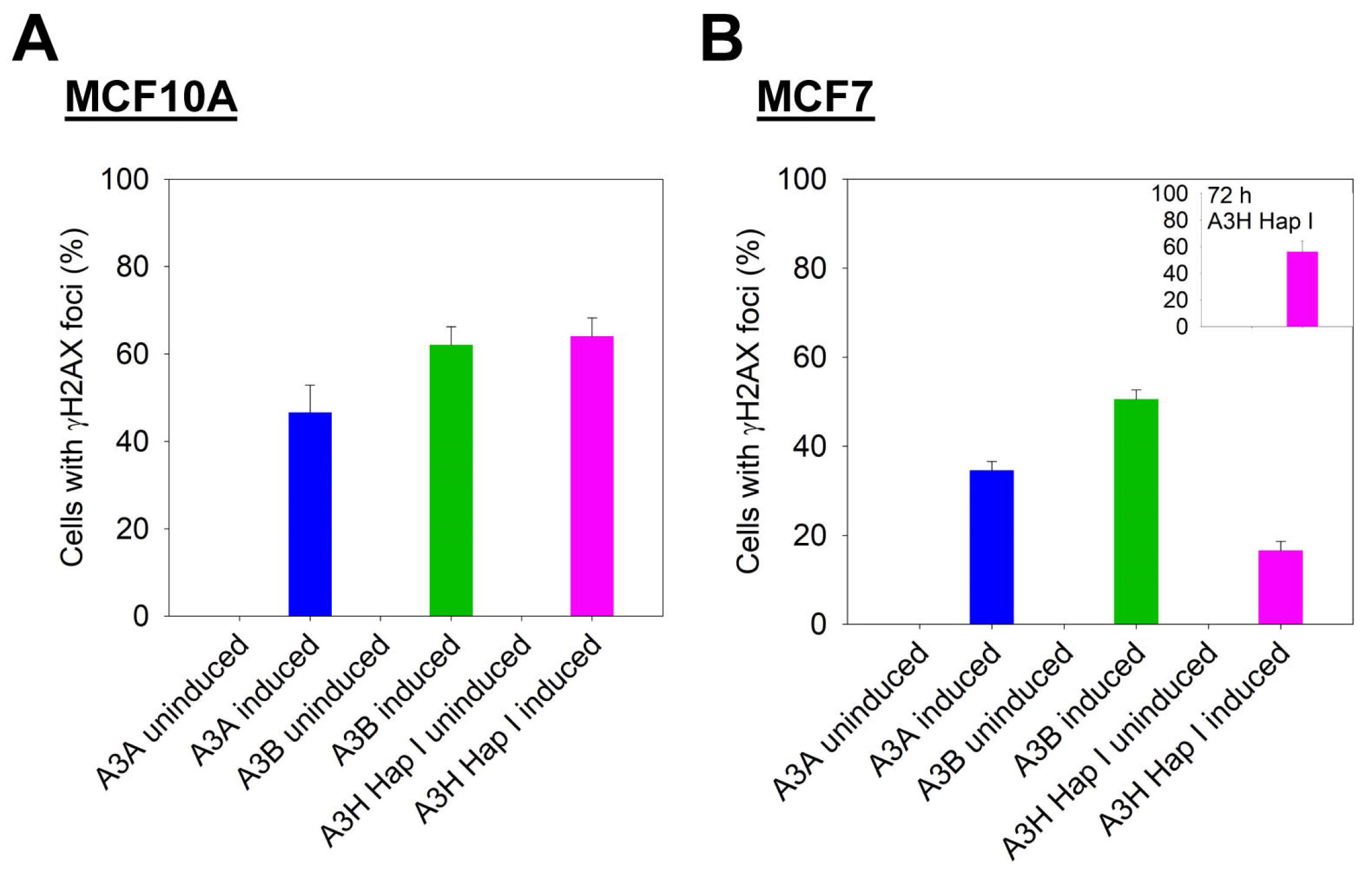
Early γH2AX foci in MCF10A and MCF7-derived stable cell lines. The A3-Flag expression was either uninduced or induced with dox for 24 h in MCF10A and MCF7 -derived stable cell lines before staining with antibodies. For A3H Hap I, the procedure was repeated in MCF7 cells after dox induction for 72 h (inset). The percentage of cells with γH2AX foci was determined. Two independent experiments were conducted. Supplementary Figures S4 and S5 show representative images from one experiment.

### 3.3 RNA in the nucleus does not inhibit A3A, A3B, or A3H Hap I cytidine deamination

Cytoplasmic RNA is known to inhibit A3B and A3H Hap I enzyme activity and this has led to the conclusion that A3A is likely more active in cells due to a lack of RNA inhibition (Cortez et al., 2019). However, the effect of RNA in the nucleus has not been studied. Based on our observation of γH2AX foci formation with all three A3s (Figures 2 and 3), we hypothesized that there is less inhibition of A3 activity in the nucleus because there is less RNA. However, before testing our hypothesis, we first wanted to reconcile the γH2AX foci formation data with previous *in vitro* data showing different A3 activities. To do this we induced or did not induce A3 expression with dox and then used WC lysates from MCF10A cells and a commonly used 43 nt substrate containing a single 5’TTC motif to test deamination activity *in vitro* (Akre et al., 2016;Serebrenik et al., 2019). Since we could not quantify the amount of A3 enzymes expressed, we quantified the extent of deamination activity as a percentage of the total substrate (Figure 4A).

**Figure 4.**
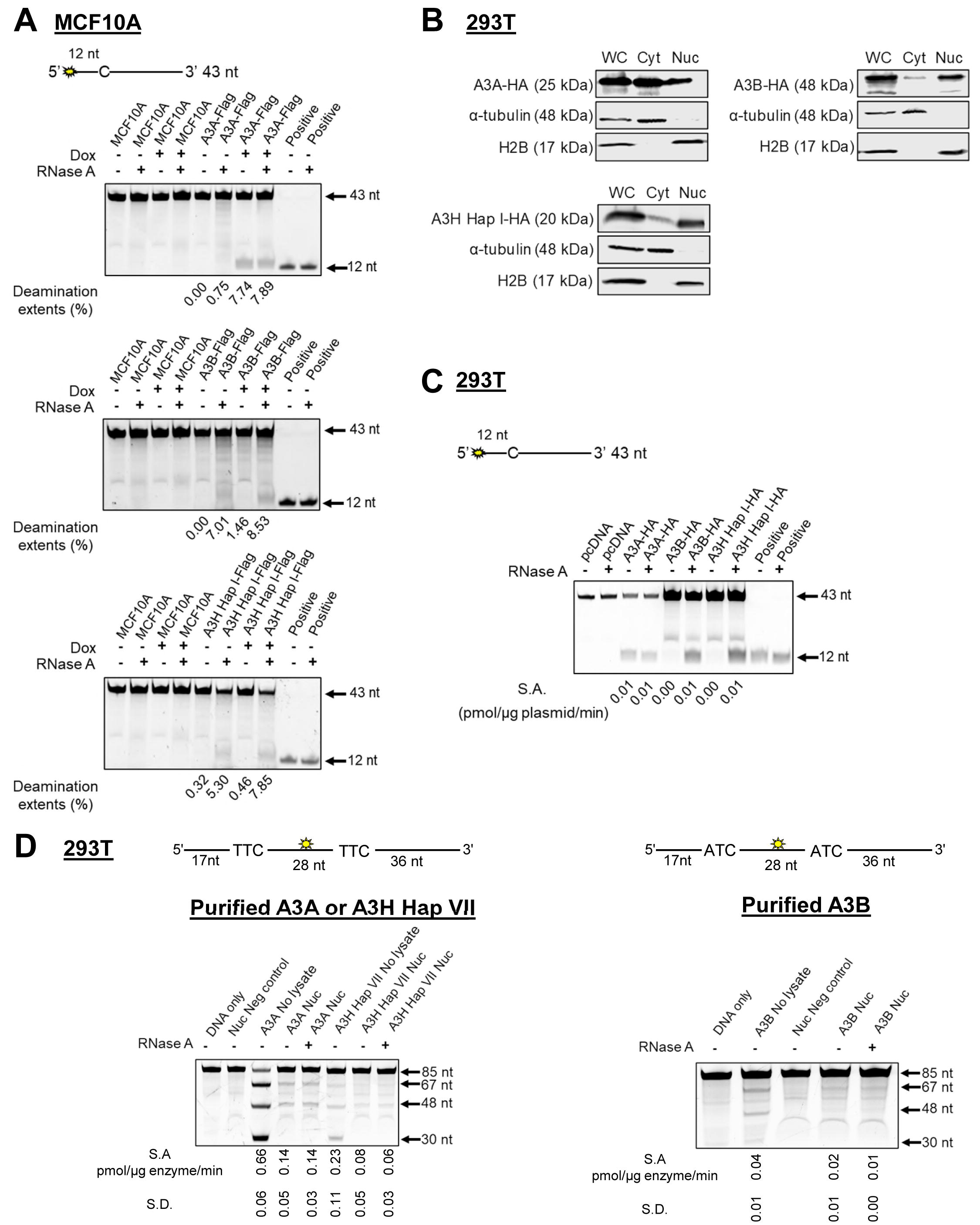
A3B and A3H are inhibited by RNA in WC lysates, but not Nuc lysates. A. Deamination assays for WC lysates from MCF10A-derived stable cell lines. The 5’-end fluorescein labeled 43 nt substrate was used at 100 nM and the reactions were carried out for 1 h (MCF10A A3A and MCF10A A3B) or 3 h (MCF10A A3H Hap I). The positive control for deamination corresponds to a 43 nt substrate containing a 5’TTU motif. The deamination extent per hour (%) is shown below the gels. **B.** 293T cells were transfected with HA-tagged A3A, A3B, or A3H Hap I and then fractionated. Protein expression and localization was assessed by immunoblot using anti-HA antibodies. The α-tubulin and H2B antibodies were used as loading controls for the Cyt and Nuc fractions, respectively. Approximate molecular weight (kDa) of each protein is shown based on the closest molecular weight marker band (Supplementary Figure S6). Blots were cropped to improve conciseness of presentation and uncropped blots are shown in Supplementary Figure S6. **C.** The activity of A3A, A3B, and A3H Hap I in transfected 293T cells was tested using a 43 nt substrate at a concentration of 50 nM for A3A and at 1 µM for A3B and A3H Hap I. The deamination reaction was carried out for 30 min (A3A), 1 h (A3B) or 2 h (A3H Hap I). The specific activity (S.A.) in WC lysates was measured as pmol substrate deaminated/µg plasmid transfected/min. **D.** Deamination activity of A3A, A3B, and A3H Hap VII in the Nuc lysates of 293T cells. Purified enzymes were added to the Nuc lysates and 85 nt substrates were used for deamination. For A3A and A3H Hap VII, 100 nM of a substrate containing two TTC motifs was used at a substrate:enzyme ratio of 1:1. For A3B, 500 nM of a substrate containing two ATC motifs was used at a substrate:enzyme ratio of 1:0.5. Reactions were carried out for 1 h and the S.A. was measured as pmol substrate deaminated/µg enzyme/min. Three biologically independent experiments were conducted and the standard deviation (S.D.) is shown. The sizes of substrate DNA and cleaved product DNA are denoted for each gel. A deamination of the 5’-proximal C results in a 67 nt band and deamination of the 3’-proximal C results in a 48 nt band, both of which are visible under all conditions. The deamination of both C residues results in a 30 nt band, but this is only visible in the No lysate condition.

Consistent with previously published results in other cell lines, in MCF10A WC lysates A3A cytidine deaminase activity was not affected by addition of RNase A, but activity was completely abolished for both A3B and A3H Hap I if the WC lysates were not treated with RNase A (Figure 4A) (Smith, 2017;Xiao et al., 2017;Feng et al., 2018;Shaban et al., 2018). Notably, in MCF10A A3B and MCF10A A3H Hap I stable cell lines, there was background deamination activity in the presence of RNase A, but no dox induction, suggesting leaky expression (Figures 1D and 4A). There was no background deamination in non-transduced parental cell lines or in the MCF10A A3A cell line suggesting that this is only specific to MCF10A A3B and MCF10A A3H Hap I cell lines. In the presence of RNase A, the *in vitro* deamination extents of A3A, A3B, and A3H Hap I were approximately equal (Figure 4A). The dox induction only increased deamination of MCF10A A3B and MCF10A A3H Hap I cell lines by ∼2% over the background, however, using immunofluorescence, we observed no background deamination activity (Figure 2B-D and Figure 4A). These data suggest that *in vitro* deamination activity from WC lysates is not an accurate estimation for in cell A3 activity.

To determine if the differences in RNA sensitivity remained the same in Nuc and Cyt lysates we sought to test fractionated cell lysates. However, we found that there was differences in the amounts of each A3 in the Nuc and Cyt lysates in 293T, MCF10A, and MCF7 cells, likely due to each A3 having a different mechanism of entering the nuclear compartment (Figure 4B and Supplementary Figure S7). Interestingly, A3A was detected in the nuclear lysates of 293T cells, but not MCF10A or MCF7 cells (Figure 4B and Supplementary Figure S7). In addition, although A3A was present in the Nuc lysates of 293T cells, the amount present for each extraction was variable. This may be because small proteins can diffuse out of the nucleus during membrane permeabilization (Ogawa and Imamoto, 2021). Although A3A and A3H Hap I have the same molecular weight, A3A is a monomer in solution (although can be a dimer when bound to DNA) and A3H Hap I is a dimer that binds RNA, resulting in a larger cellular molecular weight for A3H Hap I (Bohn et al., 2015;Bohn et al., 2017;Salamango et al., 2018). Due to the background deamination in MCF10A cells (Figure 4A), we determined if 293T cells would be suitable to test our hypothesis. We transfected a known amount of A3- expression plasmid and calculated a specific activity based on the pmol of substrate deaminated per µg of plasmid transfected per minute (Figure 4C). Similar results to MCF10A cells for deamination activity were observed with transient transfection of A3 enzymes in 293T cells, but without the background deaminase activity (Figure 4A, C).

Thus, we used Nuc lysates purified from untransfected 293T cells and added purified A3A, A3B, and A3H Hap VII to the lysates. We then calculated the specific activity based on the pmol of substrate deaminated per µg of enzyme per minute (Figure 4D). The A3H Hap VII is a stable proxy for A3H Hap I that differs only by one amino acid can be studied *in vitro* (Adolph et al., 2017b). Longer 85 nt substrates containing the preferred 5’ATC (A3B) or 5’TTC (A3A and A3H) motifs were used to ensure optimal deamination conditions. While A3A prefers hairpin substrates, we did not use a hairpin substrate for comparison between the three enzymes since A3B and A3H do not deaminate hairpin substrates efficiently (Supplementary Figure S8). Equal volumes of the same Nuc lysates were used for each biological experiment for A3A, A3B, and A3H Hap VII, which ensured that RNA and other nuclear proteins and/or factors were at the same level in each reaction. A3A was able to deaminate the DNA in Nuc lysates treated or not with RNase A, although the activity in the absence of RNase A was approximately 5-fold less than with no lysate (Figure 4D, left panel). Similar results were found on the hairpin substrate (Supplementary Figure S8). In contrast to the WC lysate data, there was no requirement for RNase A for deamination activity in Nuc lysates for A3H Hap VII (Figure 4D, left panel) and A3B (Figure 4D, right panel), although the activity was approximately 2- or 7- fold less than A3A, respectively. The A3H Hap VII and A3B activities were 3-fold and 2-fold less in the Nuc lysate than no lysate in the absence of RNase A, respectively (Figure 4D). Nonetheless, these data corroborate our hypothesis that in the nucleus, compared to WC lysates, there is less inhibition by RNA for A3B and A3H Hap VII.

Another observation specific to the Nuc lysate is that A3B and A3H Hap VII, which are processive enzymes that usually have the ability to deaminate both cytosines on the same DNA, resulting in a 30 nt band, were unable to do so under the single hit conditions of the experiment (Feng et al., 2015;Adolph et al., 2017b). Single hit conditions result when less than 15% of the substrate is used in the reaction and enables conclusions to be made about single ssDNA enzyme encounters (Creighton et al., 1995). The A3A is not processive and a 30 nt band was not expected under single hit conditions, but is visible once a large amount of substrate has been used in the reaction (see No lysate condition, Figure 4D) (Love et al., 2012).

Overall, these results considered with the γH2AX foci formation suggest that the *in vitro* data cannot be used alone to gauge activity in cells since A3A, A3B, and A3H Hap I showed different activities *in vitro* (Figure 4D), but similar activities in cells as measured by γH2AX foci (Figures 2 and 3). However, since γH2AX foci marks both dsDNA breaks and replication fork slowing, it is not known, for example, if A3A causes more DNA breaks and A3H and A3B cause only replication fork stalling due to less deaminase activity (Rogakou et al., 1998;Ward and Chen, 2001). This would be consistent with A3A being identified to cause a higher mutations frequency in cancer cells and cell lines (Petljak et al., 2022). To investigate the specific effects of each A3, we determined the ability to induce common cancer phenotypes in both MCF10A and MCF7 cells. Since MCF10A is more proficient in DNA repair than MCF7 cells, especially for double strand break repair (Francisco et al., 2008), we expected to observe cell type specific differences.

### 3.4 A3A and A3B affect anchorage independent growth in soft agar

The soft agar colony formation assay is used to measure anchorage independent growth, a hallmark of cellular transformation. We compared each cell line exposed to dox to its dox untreated condition. In the non-tumorigenic model cell line MCF10A, no differences were observed between the dox treated/untreated condition (∼15 colonies) (Figure 5A-B). However, significant differences were observed for A3A expressing cells that showed a decrease in colony formation from ∼20 to ∼10 colonies (Figure 5A-B). There were no significant differences observed for colonies exposed to A3B and A3H Hap I under normal replication conditions in MCF10A cells. Additionally, we tested if the effects of A3-mediated DNA damage would be amplified during replication fork stalling. Cells were either untreated or treated with dox for 24 h and subsequently treated with 2 mM HU for 6 h to induce fork stalling before plating in soft agar. Similar to the normal replication fork conditions, we did not observe differences in the number of colonies between uninduced/induced parental or A3H Hap I (Figure 5A-B). However, statistically significant differences were observed between uninduced/induced A3A and A3B. The uninduced +HU A3A condition had ∼35 colonies/field in comparison to the induced +HU A3A condition which had ∼20 colonies/field. Similarly, uninduced +HU A3B condition had ∼25 colonies/field in comparison to the induced +HU A3B condition which had <10 colonies/field. These data show that A3A and A3B have deleterious effects on soft agar colony formation in the presence of replication stress and A3A has this effect even under normal replication conditions.

**Figure 5.**
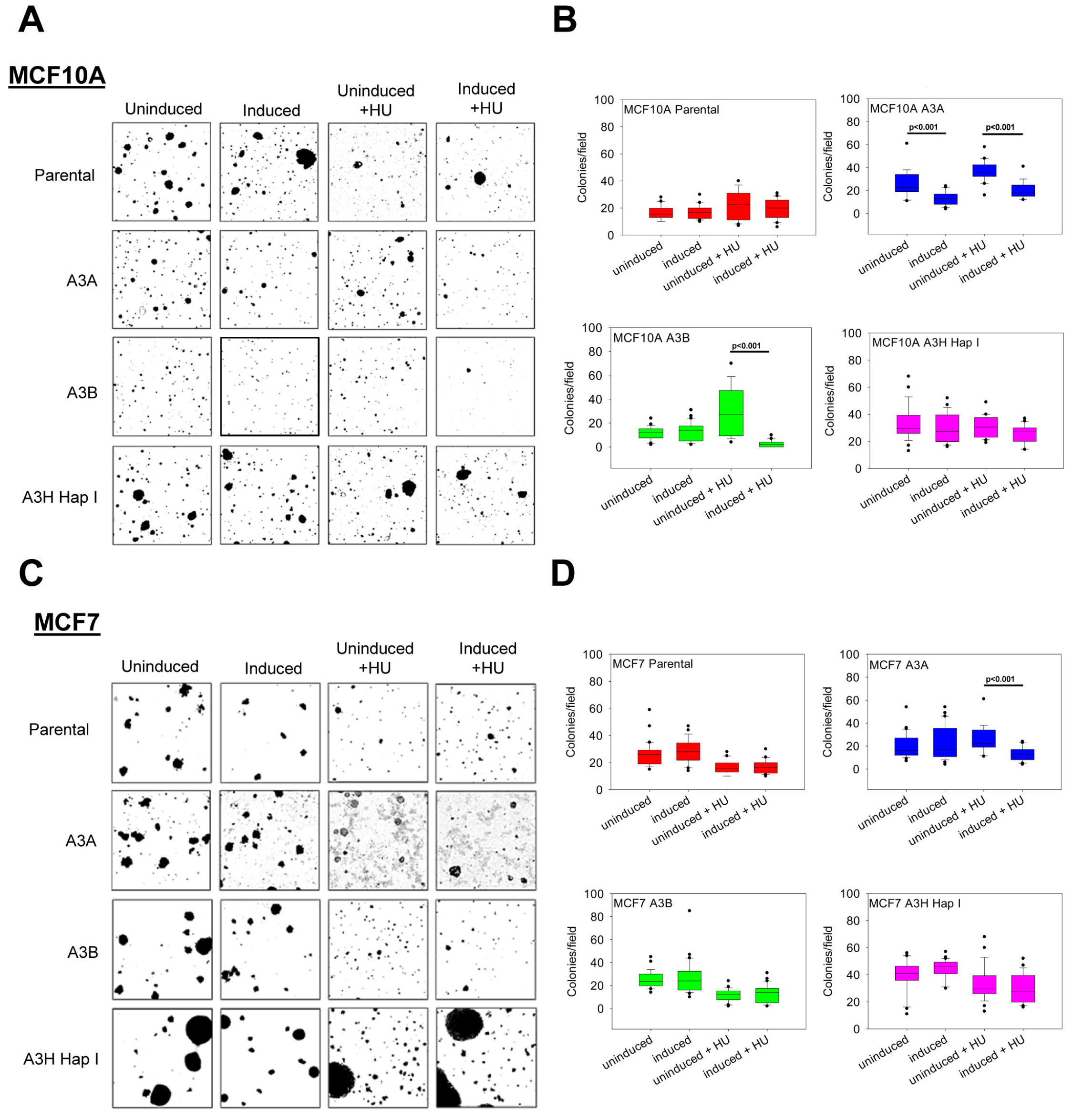
Effect of the expression of A3A, A3B, and A3H Hap I on anchorage independent growth in soft agar. MCF10A and MCF7 cells were either untreated, treated with dox (0.05- 2 μg/mL, 24 h) or treated with dox and HU (2 mM-12 mM, 6 h). After 3 weeks, colonies were imaged and quantified. **A.** Representative images of colony formation in uninduced, induced, uninduced+ HU, and induced+ HU MCF10A, A3A, A3B and A3H Hap I cells. **B.** Box-whisker plot of quantified uninduced, induced, uninduced +HU, and induced +HU MCF10A colonies. **C.** Representative images of colony formation in uninduced, induced, uninduced +HU, and induced +HU MCF7, A3A, A3B and A3H Hap I cells. **D.** Box-whisker plot of quantified uninduced, induced, uninduced +HU and induced +HU MCF7 colonies. Three biologically independent experiments were conducted. One representative experiment is shown in panels A and C. The statistical differences were determined by a t test on the biologically independent experiments and considered statistically significant for p<0.05.

We also tested the tumorigenic MCF7 breast cancer cell lines and did not observe differences in the number of colonies between the uninduced and induced condition for parental cells or cells that expressed either A3A, A3B or A3H Hap I (Figure 5C-D). During replication fork stalling using HU treatment no difference in colony growth was observed between uninduced +HU and induced +HU, for A3B or A3H Hap I. A3H Hap I colonies grew larger than A3B or A3A colonies, but this did not affect the total number of colonies. A statistically significant difference was observed in the A3A condition where uninduced +HU colonies were significantly greater (∼25 colonies/field) than in the induced +HU condition (∼10 colonies/field). These data show that A3A expression is more deleterious to MCF7 cells when combined with replication stress (Figure 5C-D).

### 3.5 A3H Hap I can induce the migration of MCF7 cells

We also investigated if the expression of A3A, A3B and A3H Hap I either in MCF10A or MCF7 cells could influence cell migration in the absence or presence of HU. Induction of A3s with or without HU treatment, had no effect in the non-tumorigenic MCF10A cells (Figure 6A). The expression of A3A and A3B either in MCF7 breast cancer cells treated or not with HU did not show any effect on cell migration (Figure 6B). However, for MCF7 cells, the expression of A3H Hap I in the presence of HU promoted cell migration compared to the same cells expressing A3H Hap I without HU and to the uninduced condition with or without HU (Figure 6B).

**Figure 6.**
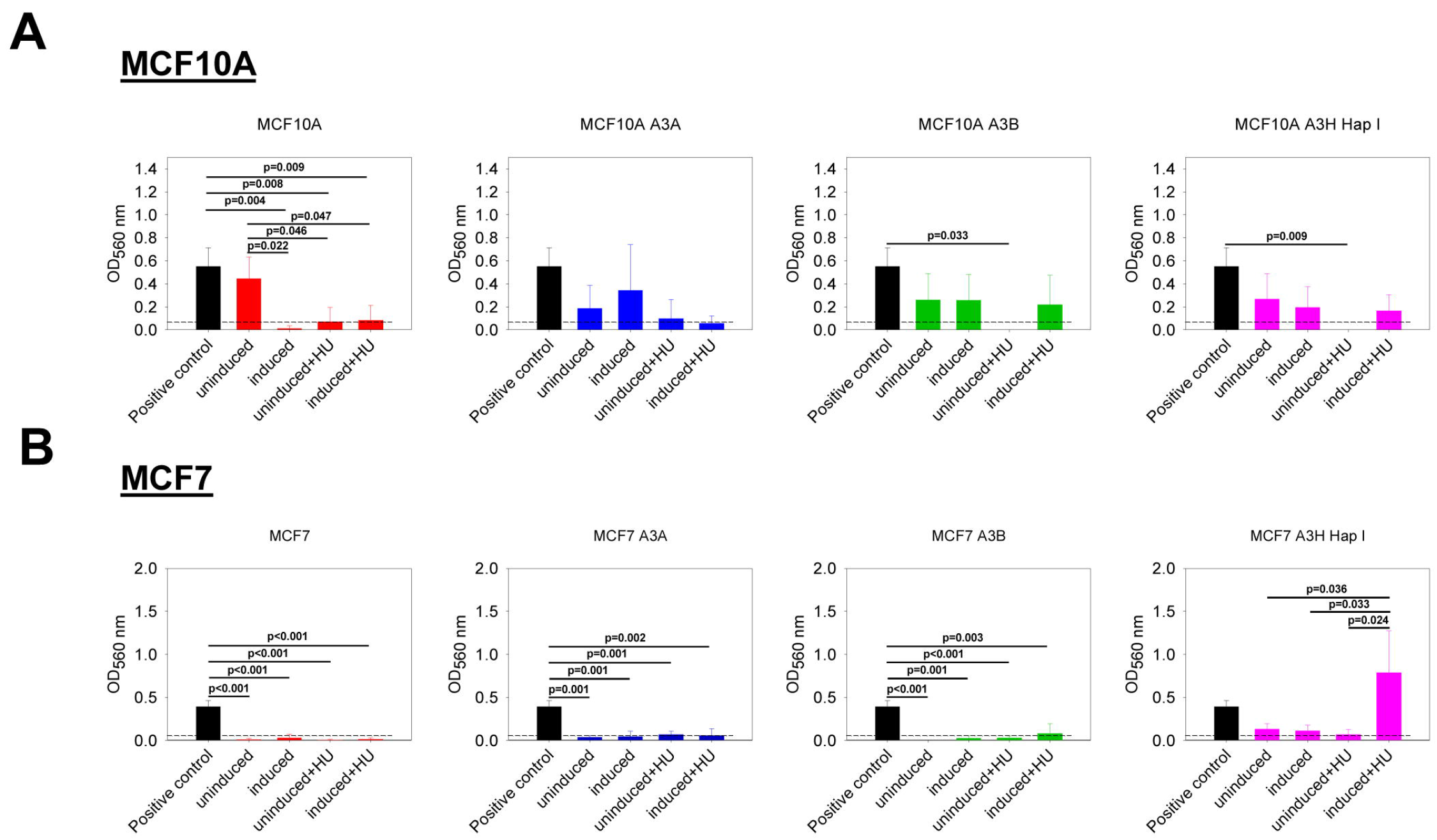
Effect of the expression of A3A, A3B, and A3H Hap I on cell migration. A colorimetric kit based on the Boyden Chamber principle was used to assess transwell migration. The OD values were measured at 560 nm. The dashed line represents the cut off calculated from the condition without cells. The BC cell line HCC1428 was used as a positive control of migration (Veeraraghavan et al., 2014). **A.** MCF10A-derived stable cell lines were either untreated (uninduced) or treated with dox (induced) for 7 d. The cells were also treated (induced+HU) or not (uninduced+HU) with dox for 7 d and twice with HU. **B.** MCF7-derived stable cell lines were treated as described for MCF10A-derived stable cell lines. Three biologically independent experiments were conducted and results are represented using mean values and SD. The statistical differences between groups were determined by one way ANOVA for p<0.05 and p values are shown on the graph.

## 4 Discussion

The existing data implicating A3A, A3B, and A3H Hap I in cancer have led to an unclear understanding of the relative contributions of individual A3s to mutagenesis in BC (Burns et al., 2013a;Burns et al., 2013b;Starrett et al., 2016;Cortez et al., 2019). To overcome these limitations, we developed MCF10A and MCF7 derived stable cell lines expressing dox inducible Flag-tagged A3A, A3B, and A3H Hap I and identified unique phenotypes for each A3. We demonstrated that all the expressed enzymes localize to the nucleus and are enzymatically active, causing UDG-dependent DNA damage. We found that *in vitro* deamination assays do not directly correlate with accumulation of DNA damage in cells or extent of inhibition by cellular RNA. Altogether, the data support a model in which A3A, A3B and A3H Hap I all induce DNA damage in normal or tumorigenic breast epithelial cells, but the phenotypic effects of the induced damage is unique for each A3.

While *in vitro* deamination in cell lysates is often used to gauge A3 activity in cells, we found inconsistencies with this method. Importantly, despite reports that A3A is a more potent inducer of DNA damage than A3B (Landry et al., 2011;Burns et al., 2013a;Mussil et al., 2013;Caval et al., 2014), our results showed that A3B and A3H Hap I could cause similar levels of γH2AX foci as A3A (Figures 2 and 3). To investigate how the deamination activity observed in cells relates to the *in vitro* deamination assay, we conducted deamination assays under various conditions. A3B and A3H Hap VII deaminated the ssDNA in the Nuc lysates as A3A does, either in presence or absence of RNase A (Figure 4D). This is the first direct evidence demonstrating that in the nucleus the A3 enzymes involved in deaminating genomic DNA are still active even without the addition of RNase A. This is also the first evidence that A3B and A3H Hap VII, that have been characterized *in vitro*as processive enzymes, are non-processive on ssDNA in nuclear lysates. This is a positive feature for genome editing where off-target deaminations at adjacent cytosines would be detrimental. In addition, despite A3A being more active in Nuc lysates *in vitro*, all three enzymes had equal ability to induce γH2AX foci in cells (Figures 2 and 3). Overall, our data suggest that A3A, A3B, and A3H Hap I each contribute to cytosine deamination and DNA damage and directly show that between A3A and A3B, neither is more active in cells. However, there must be other pressures in the cells that result in a more detrimental effect of A3A on anchorage independent growth of cells (Figure 5) and more A3A-induced mutations being observed in cell line experiments (Petljak et al., 2022). This will be an area of future study and may be related to the superior ability of A3A to displace RPA (Adolph et al., 2017b;Wong et al., 2021).

In this study, we looked for phenotypic markers of A3 expression. We observed that A3A expression could delay anchorage independent growth of cells in soft agar (Figure 5). A3A was most able to cause a decrease in the number of colonies formed in this assay in the presence or absence of HU in MCF10A cells or in the presence of HU in MCF7 cells (Figure 5). In contrast, A3B was only able to decrease proliferation of MCF10A cells in the presence of HU. Based on previous data which shows the tumorigenic potential of A3A (Law et al., 2020), it appears that A3A may cause lethal damage to a large population of cells, but surviving ones may have tumorigenic potential. A3A has been found to cause deamination-independent chromosomal instability in pancreatic ductal adenocarcinoma, but only if there is existing cellular alterations such as in p53 function of KRAS signaling (Wörmann et al., 2021). It remains to be determined if this occurs in other cancers, such as BC. In this study we did not find that catalytic mutants of A3A (or A3B or A3H Hap I) could cause chromosomal instability as measured by γH2AX foci in MCF10A or MCF7 cells, although other measures of chromosomal instability remain to be tested (Figure 2).

There have been few studies that have examined A3H Hap I and its effect on cancer. The A3H mutation signature is found in lung adenocarcinomas (Starrett et al., 2016). A genetic and biochemical study showed that an A3H Hap I single nucleotide polymorphism that destabilized the enzyme was beneficial for lung cancer, suggesting that A3H Hap I activity would be detrimental to cancer cell growth or increase immune recognition (Hix et al., 2020). However, the role of A3H in BC is less clear (Starrett et al., 2016). Our results showed that the MCF7 cells expressing A3H Hap I with a pre-existing replication stress induced by HU treatment were able to migrate (Figure 6). These are the first direct evidence of the potential of A3H Hap I to contribute to a cancer cell phenotype.

In summary, we demonstrated that all three A3s are able to induce cellular phenotypes as a result of their induced DNA damage. Importantly we showed that cytidine deamination activity for all A3s was not inhibited by RNA in Nuc extracts and was similar in cells, even with differing *in vitro* activities. These data demonstrate the importance of using accurate *in vitro* systems to determine cytidine deaminase activity and necessitate a reassessment of the contributions of A3A, A3B, and A3H Hap I to somatic mutagenesis during tumorigenesis. These data also provide more detailed deamination activity analysis for A3A, A3B, and A3H Hap I in the context of nuclear DNA that can be used in the design of base editing technologies.

## 6 Author Contributions

MGR and LC designed the study. MGR and LW performed experiments. MGR and LC wrote the manuscript. MGR, LW, and LC edited the manuscript.

## Acknowledgements

We acknowledge Dr. Amit Gaba for critical comments during the writing of this article, Dr. Deborah Anderson for technical advice, Aimen Khan for technical support, and Arzhang Shayeganmehr for conducting the site directed mutagenesis for the A3A and A3H Hap I catalytic mutants.

