## Supplementary Data for "Similar deamination activity but different phenotypic outcomes induced by APOBEC3 enzymes in breast epithelial cells"

*Corresponding author

**Table S1. Probe and primer sequences used for qPCR.**

| Name | Sequence |
|---|---|
| A3A Forward | 5‘ TCT CCA TCA TGA CCT ACG ATG A |
| A3A Reverse | 5‘ CCT GGT GGT CCA CAA AGG T |
| A3A Probe | 5‘ CTG CTG GG |
| A3H Forward | 5‘ AGC CGA AAC ATT CCG CTT AC |
| A3H Reverse | 5‘ CGT CAG CTG GTA ACA CAA GAG |
| A3H Probe | 5‘ CTG AGG CGG CGC TTG TTG TTA AAC |

The A3A probe was purchased from Roche as Universal Probe Library #3 probe. The A3H probe was designed to contain a 5' fluorescein, an internal Zen Quencher, and a 3‘Iowa Black FQ Quencher.

**Table S2. Site directed mutagenesis primers.**

| Name | Sequence |
|---|---|
| A3A E72Q Forward | 5’ TAC GGC CGC CAT GCG CAG CTG CGC TTC TTG GAC |
| A3A E72Q Reverse | 5’ GTC CAA GAA GCG CAG CTG CGC ATG GCG GCC GTA |
| A3B E255Q Forward | 5’ GCC GCC ATG CGC AGC TGC GCT TC |
| A3B E255Q Reverse | 5’ GAA GCG CAG CTG CGC ATG GCG GC |
| A3H E56Q Forward | 5’ GAA AAA GTG CCA TGC ACA AAT TTG CTT TAT TAA C |
| A3H E56Q Reverse | 5’ GTT AAT AAA GCA AAT TTG TGC ATG GCA CTT TTT C |

**Table S3. ssDNA substrates used for deamination assays.**

| Name | Sequence |
|---|---|
| 43 nt TTC | 5' fluorescein ATTATTATTATTCGAATGGATTTATTTATTTATTTATTTATTT |
| 43 nt TTU | 5' fluorescein ATTATTATTATTUGAATGGATTTATTTATTTATTTATTTATTT |
| 85 nt ATC with deamination motifs 30 nt apart | 5' AAA GAG AAA GAG TAA ATC AAA GAG TAA AGT {dT-FAM} AAG TAG AGA GAT TAT ATC AAA GAG TAA AGT TAG TAA GAT GTG TAA GTA TGT TAA |
| 85 nt TTC with deamination motifs 30 nt apart | 5' AAA GAG AAA GAG TAA TTC AAA GAG TAA AGT {dT-FAM} AAG TAG AGA GAT TAT TTC AAA GAG TAA AGT TAG TAA GAT GTG TAA GTA TGT TAA |
| 21 nt TTC hairpin | GCA AGC TGT TCA GCT TGC TGA |
| 21 nt ATC hairpin | GCA AGC TGA TCA GCT TGC TGA |

The deamination motifs are underlined.

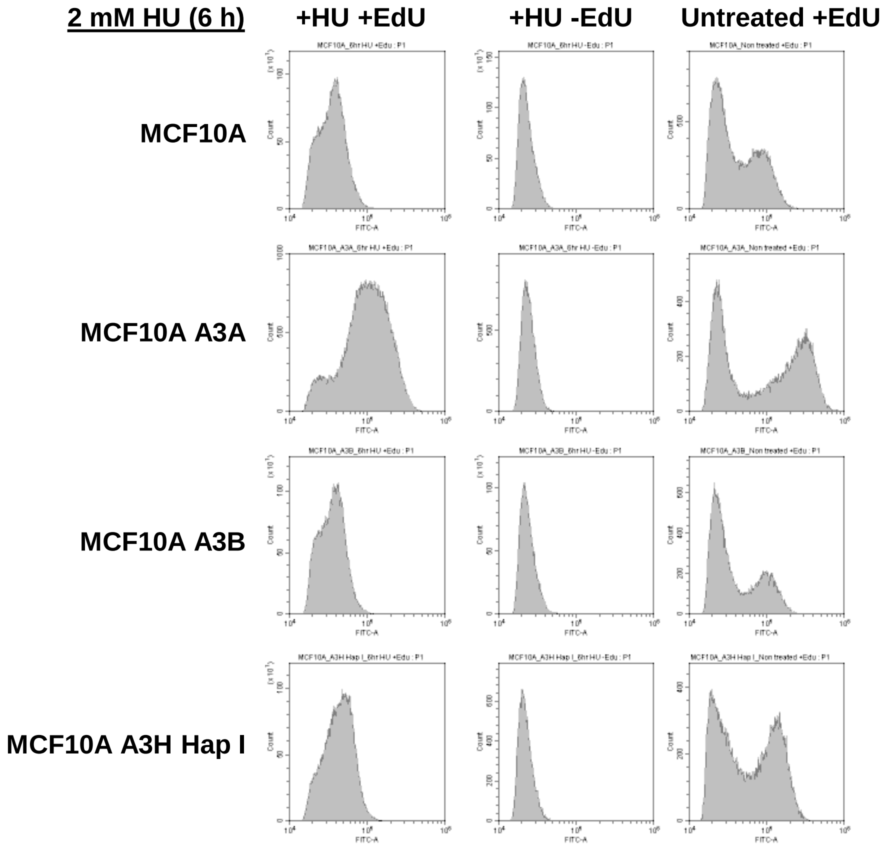


**Supplementary Figure S1. Evaluation of HU treatment to stall the replication fork for MCF10A-derived stable cell lines.** Cells were treated with 2 mM HU for 6 h and then processed using Click-iT® EdU Flow Cytometry Assay Kit from Molecular Probes according to the manufacturer's instructions. To detect proliferating cells which have incorporated EdU and nonproliferating cells which have not, cells were labeled with Alexa Fluor® 488 azide and analyzed by flow cytometry on a Cytoflex Cytometer using 488 nm excitation with a green emission filter.

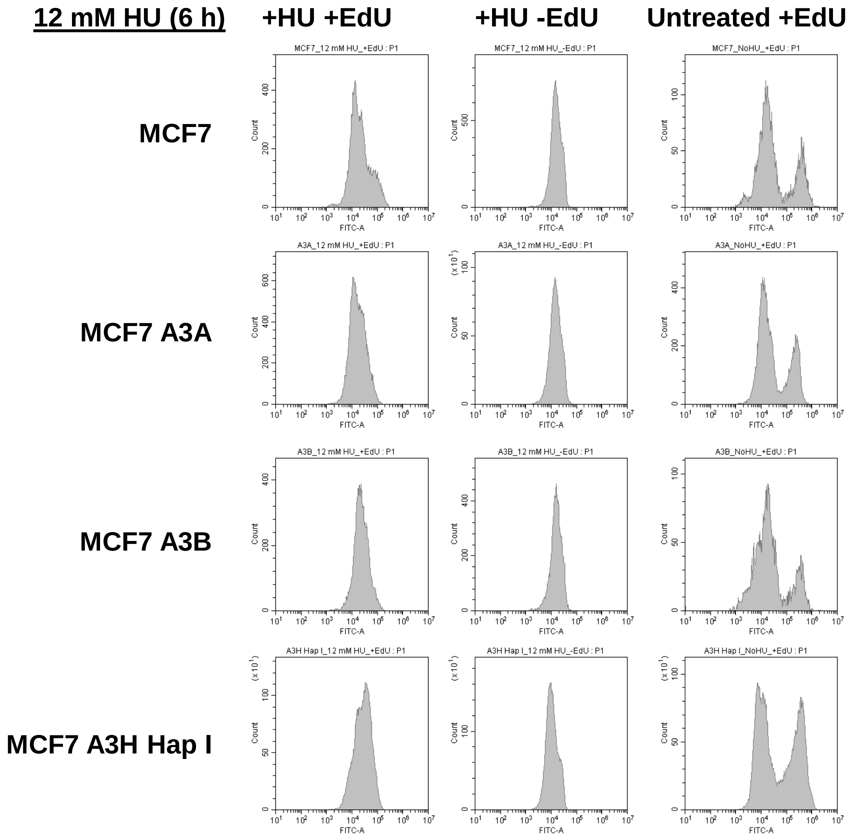


**Supplementary Figure S2. Evaluation of HU treatment to stall the replication fork for MCF7-derived stable cell lines.** Cells were treated with 12 mM HU for 6 h and then processed using Click-iT® EdU Flow Cytometry Assay Kit from Molecular Probes according to the manufacturer's instructions. To detect proliferating cells which have incorporated EdU and nonproliferating cells which have not, cells were labeled with Alexa Fluor® 488 azide and analyzed by flow cytometry on a Cytoflex Cytometer using 488 nm excitation with a green emission filter.

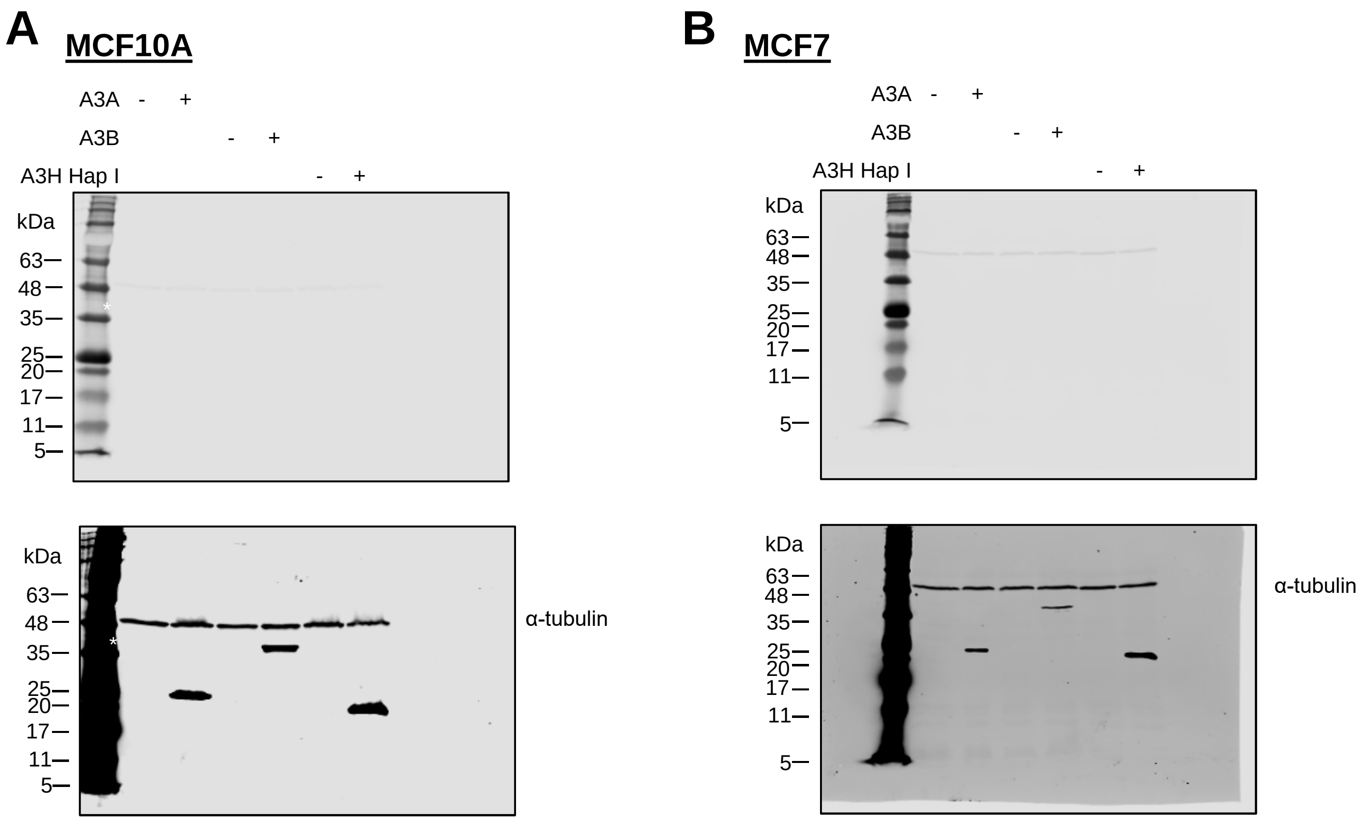


**Supplementary Figure S3. Immunoblots used in Figure 1B-C.** A. Expression of A3A, A3B, and A3H Hap I in MCF10A cells without (-) or with (+) the addition of dox. B. Expression of A3A, A3B, and A3H Hap I in MCF7 cells without (-) or with (+) the addition of dox. The α-tubulin loading control was visualized on the same blot as the A3-Flag but using a different secondary antibody. Upper and lower panel are low and high contrast versions of the same blot, respectively, to enable visualization of individual bands of the molecular weight marker.

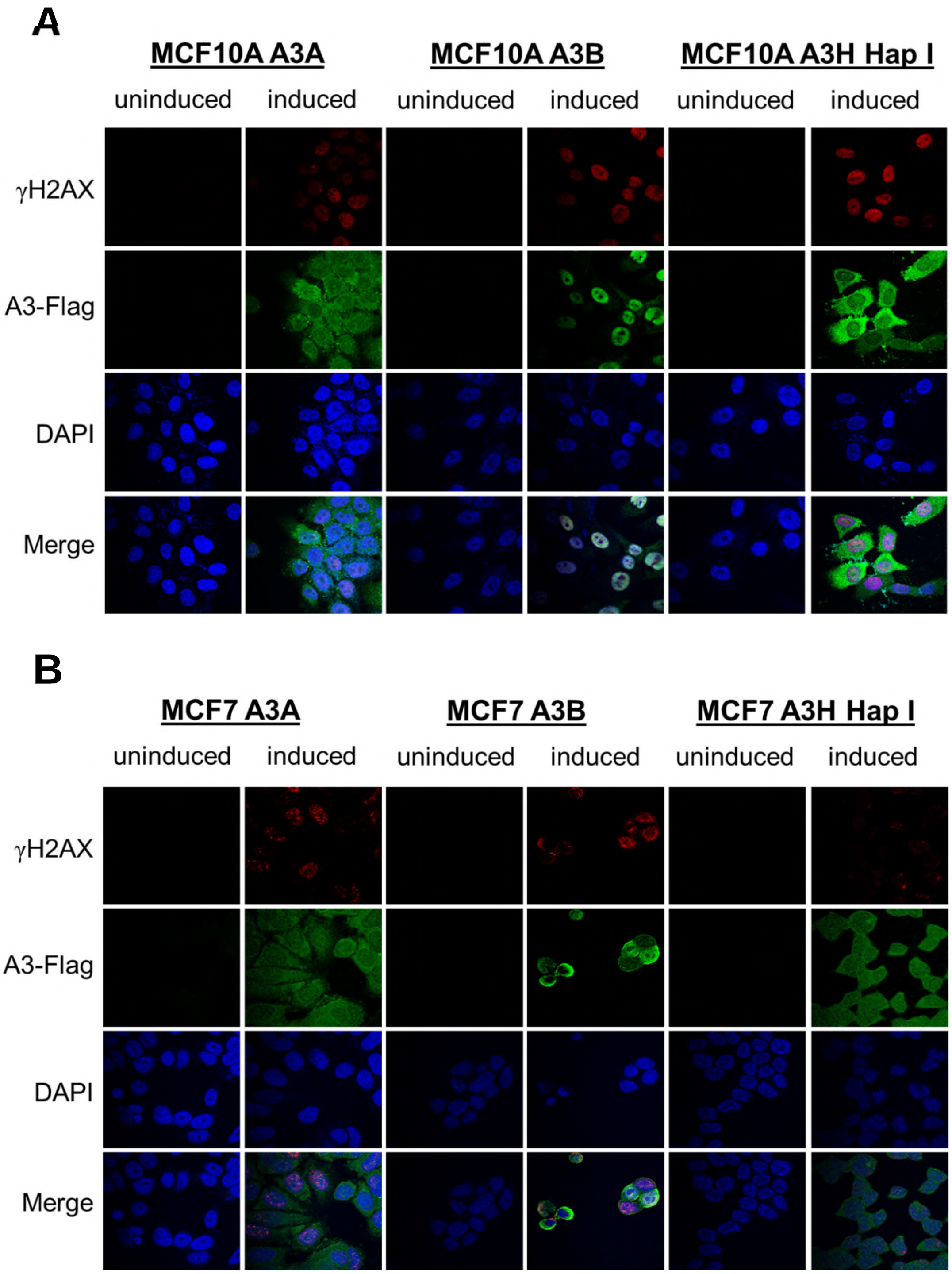


**Supplementary Figure S4. Detection of A3-induced γH2AX foci formation in MCF10A- and MCF7- derived stable cell lines after 24 h.** The Flag-A3 expression was either uninduced or induced with dox in (A) MCF10A- or (B) MCF7- derived stable cell lines before staining with antibodies. Green color identifies Flag-tagged A3 proteins; red color indicates γH2AX foci formation, and blue color indicates nuclei stained with DAPI.

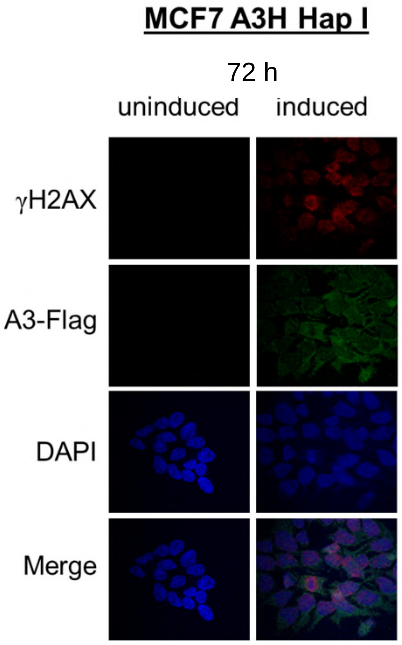


**Supplementary Figure S5. Detection of A3H Hap I-induced γH2AX foci formation in MCF7 derived stable cell line after 72 h.** The Flag-A3 expression was either uninduced or induced with dox in the MCF7-A3H Hap I stable cell lines before staining with antibodies. Green color identifies Flag-tagged A3 proteins; red color indicates γH2AX foci formation, and blue color indicates nuclei stained with DAPI.

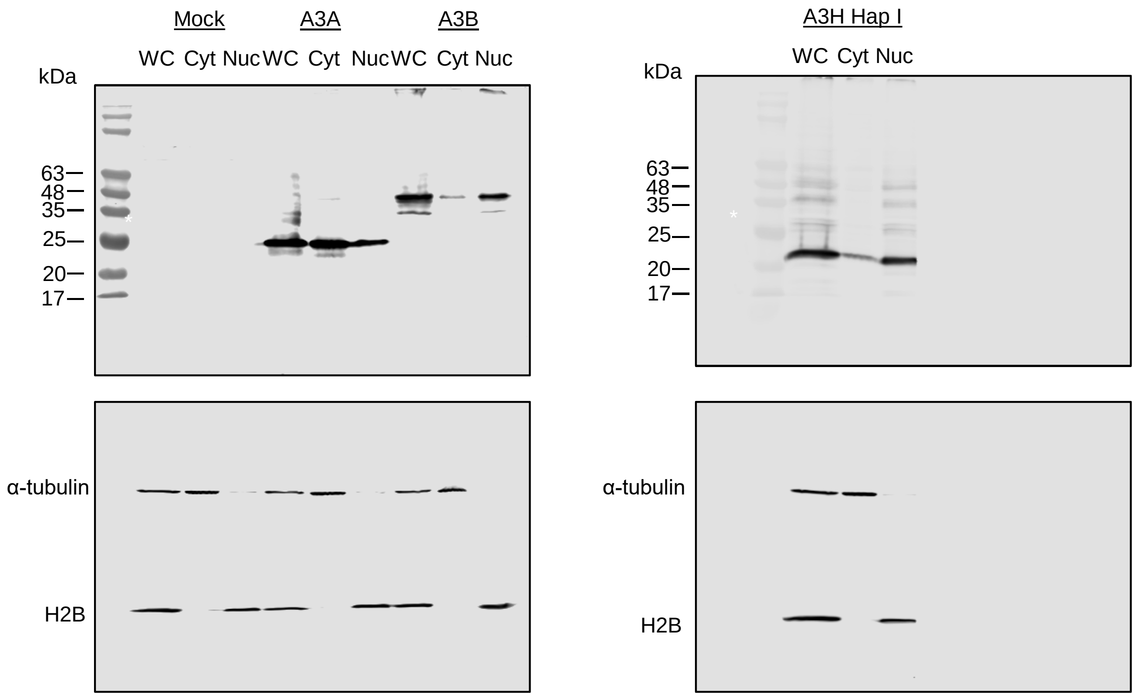


**Supplementary Figure S6. Immunoblots used in Figure 4B.** Cropped images in Figure 4B were taken from these blots. The α-tubulin and H2B loading controls were visualized on the same blot as the A3-Flag but using a different secondary antibody.

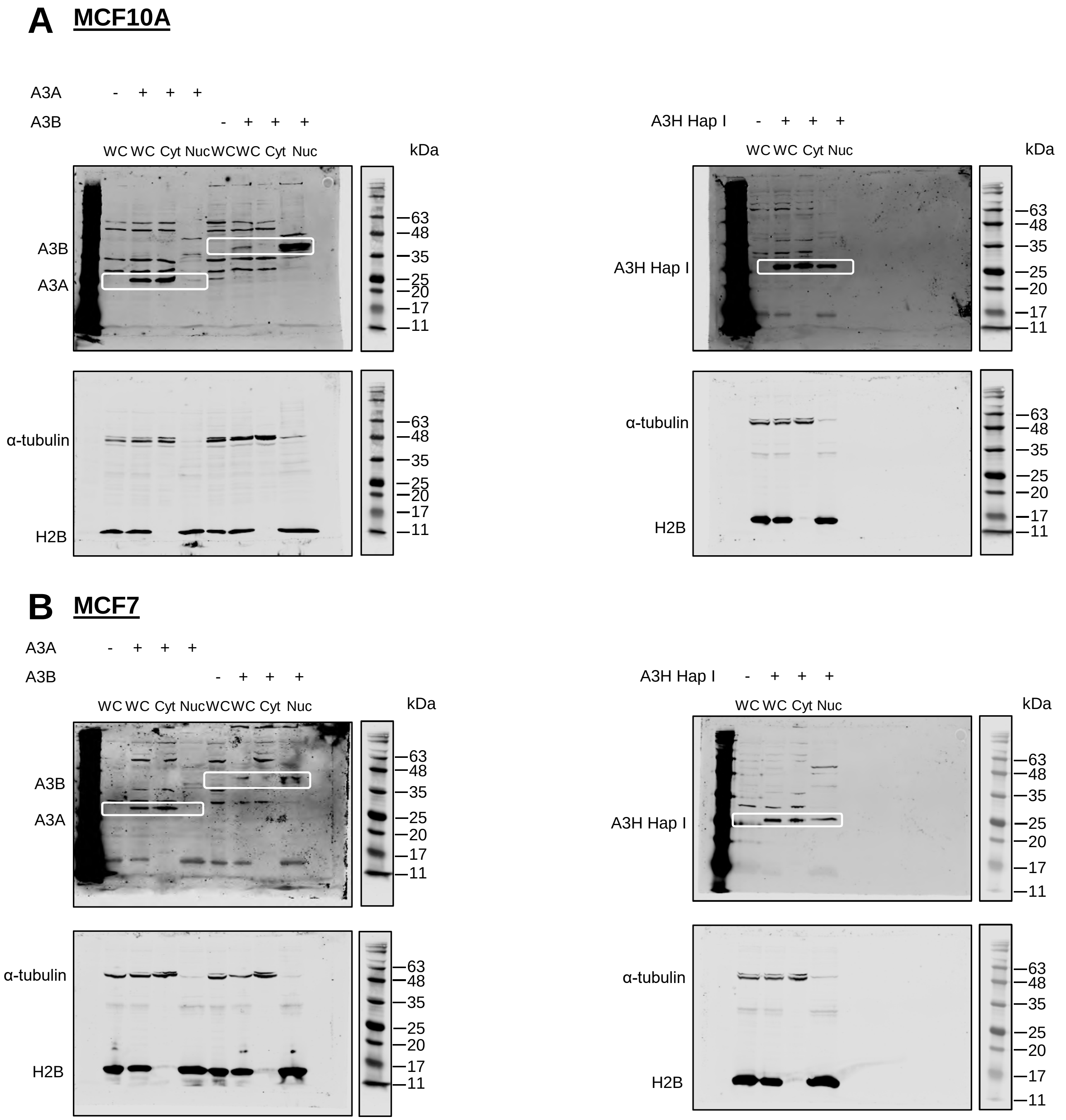


**Supplementary Figure S7. Protein localization assessed by cell fractionation for MCF10A and MCF7-derived cell lines.** A-B. MCF10A- and MCF7- derived stable cell lines were untreated (-) or treated (+) with dox for 72 h and then fractionated. Protein localization in WC, Cyt and Nuc lysates was assessed using anti-Flag antibodies. The α-tubulin and H2B antibodies were used as loading controls for the Cyt and Nuc fractions, respectively**.** The α-tubulin and H2B were detected on the same blot as the A3-Flag, but using a different secondary antibody. The molecular weight marker from a low contrast version of the same blot is shown to the right. Three independent experiments were conducted and one representative blot is shown.

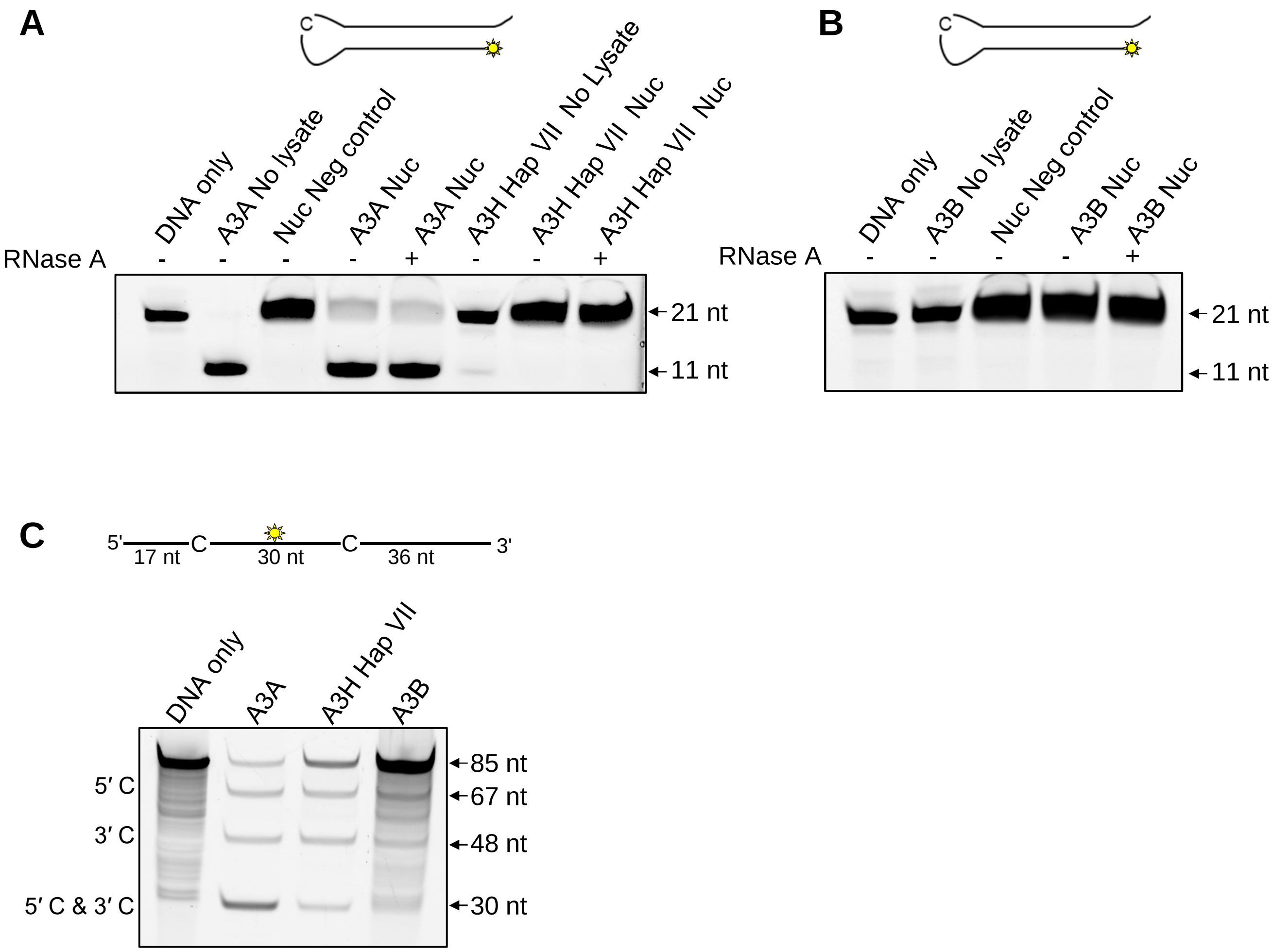


**Supplementary Figure S8. A3A, but not A3H and A3B are active on hairpin substrates.** Deamination activity of A3A, A3B, and A3H Hap VII in the Nuc lysates of 293T cells. Purified enzymes were added to the Nuc lysates and a 21 nt hairpin substrate (S3 Table) was used for deamination. **A.** The 5'TTC substrate was used for A3A and A3H Hap VII. **B.** The 5'ATC substrate was used for A3B. For A3A and A3H Hap VII, 100 nM of substrate was used at a substrate:enzyme ratio of 1:1. For A3B, 500 nM of substrate was used at a substrate:enzyme ratio of 1:0.5. The reactions were carried out for 1 h. **C.** Deamination activity on a linear 85 nt substrate (either 5'TTC for A3A and A3H Hap VII or 5'ATC for A3B) was assessed in parallel to the deamination activity on the hairpin substrate to confirm that A3H Hap VII and A3B were active in the 293T Nuc lysates. The sizes of substrate DNA and cleaved product DNA are denoted for each gel. The experiments were performed in duplicate and representative images are shown.
